# Subarachnoid hemorrhage leads to early and persistent functional connectivity and behavioral changes in mice

**DOI:** 10.1101/826891

**Authors:** David Y Chung, Fumiaki Oka, Gina Jin, Andrea Harriott, Sreekanth Kura, Sanem Aykan, Tao Qin, William J Edmiston, Hang Lee, Mohammad A Yaseen, Sava Sakadžić, David A Boas, Michael J Whalen, Cenk Ayata

## Abstract

Aneurysmal subarachnoid hemorrhage (SAH) leads to significant long-term cognitive deficits. Studies in survivors of SAH show an association between persistent cognitive deficits and alterations in resting state functional connectivity (RSFC). However, modalities commonly used to assess RSFC in humans, such as fMRI, have practical limitations in small animals. Therefore, we used non-invasive functional optical intrinsic signal imaging to determine the effect of SAH on measures of RSFC in mice at early (day 4), intermediate (1 month), and late (3 months) time points after prechiasmatic arterial blood injection. We assessed Morris water maze, open field test, Y-maze, and rotarod performance from approximately 2 weeks to 3 months after SAH induction. We found qualitative and quantitative differences in seed-based connectivity maps between sham and SAH mice. SAH reduced motor, retrosplenial and visual seed-based connectivity indices, which persisted in retrosplenial and visual cortex seeds at 3 months. Seed-to-seed connectivity analysis confirmed attenuation of correlation coefficients in SAH mice, which persisted in predominantly posterior network connections at later time points. Seed-independent global and interhemispheric indices of connectivity revealed decreased correlations following SAH for at least 1 month. SAH led to Morris water maze hidden platform and open field deficits at 2 weeks, and Y-maze deficits for at least 3 months, without altering rotarod performance. In conclusion, experimental SAH leads to early and persistent alterations both in hemodynamically-derived measures of RSFC and in cognitive performance.

## INTRODUCTION

Subarachnoid hemorrhage (SAH) from a ruptured brain aneurysm is a severe form of stroke which leads to significant mortality and long-term functional deficits [1, 2]. Many survivors of SAH suffer from persistent cognitive problems [3] with the majority not being able to return to work or resume their previous activities [4]. Recent studies using functional neuroimaging have found an association between these persistent cognitive deficits and alterations in neuronal networks after SAH using fMRI or magnetoencephalography [5–7]. In one study, altered measures of resting state functional connectivity (RSFC) were associated with worse performance on the Montreal Cognitive Assessment, a test of mild cognitive impairment [7]. Another study found that altered RSFC was associated with decreased executive dysfunction and worse quality of life as measured by a battery of standard cognitive tests [6]. However, a causal relationship between altered functional connectivity and cognitive deficits is not yet established, in part because of a lack of experimental models.

Therefore, we sought to determine if SAH causes changes in functional connectivity and associated cognitive deficits longitudinally in a mouse model of SAH. Because there are significant practical challenges in performing fMRI in mice—such as low resolution, magnet availability, and maintenance of anesthesia while in the scanner—we examined RSFC using non-invasive optical intrinsic signal imaging which detects changes in blood volume analogous to the fMRI BOLD signal.

## METHODS

### Animals

The study was carried out adhering to ARRIVE guidelines, including blinding [8]. All experiments were performed according to the Guide for Care and Use of Laboratory Animals (NIH Publication No. 85-23,1996) and protocols approved by the Institutional Animal Care and Use Committee (MGH Subcommittee on Research Animal Care). Male C57BL/6J mice (Charles River Laboratories, Wilmington, MA, USA) between 10-20 weeks of age were used for all experiments. Two animals died during imaging sessions as a result of anesthetic overdose, one each at intermediate and late time points). One animal was excluded from the study due to barbering behavior.

### SAH induction and imaging window placement

For induction of SAH, we used the anterior prechiasmatic injection approach [9, 10]. Mice were anesthetized with isoflurane (5% induction, 1-1.5% maintenance) in a mixture of 70:30 N_2_O:O_2_ and allowed to breathe spontaneously. After fixing the head in a stereotaxic frame, the scalp was incised to expose the dorsal skull surface. At this stage, an RFID glass capsule (sparkfun.com, SEN-09416) was inserted into the dorsal cervical area through the scalp incision. A 0.7 mm burr hole was drilled 5 mm anterior to bregma and slightly off of the midline to avoid the sagittal sinus. Non-heparinized blood was freshly collected from the femoral artery of a littermate donor. A 27G Whiteacre spinal needle (BD ref # 405079) connected to a 1 cc syringe was positioned with the side port facing up and angled 35° from the vertical axis. The needle was advanced carefully until the base of the skull was contacted (approximately 7 mm in most animals) then the needle was retracted 0.5-1.0 mm. Blood was injected (80 μl) over 10 seconds using a digital infusion pump. The needle was kept in place for an additional 2 min and then removed. For sham animals, the procedure was identical—including insertion of the needle and the duration of each step--except that there was no blood injection; saline injection was not done to avoid intracranial pressure increase, an important mechanism of injury in SAH.

Following SAH or sham procedure, a chronic glass coverslip was placed as previously described [11]. In brief, the surface of the skull was cleared of any residual blood products or debris. A glass coverslip was cut to approximate the shape of the dorsal skull surface. C&B Metabond cement was mixed using Clear L-Powder (Parkell, Edgewood, NY, USA) and used to adhere the glass coverslip to the exposed skull. Additional cement was applied to fill any remaining gaps between the skull and coverslip. The cement was cured for an additional 15 minutes and the animal allowed to recover.

### Imaging

Following SAH induction or sham surgery, RSFC was performed at early (post-operative day 4 with the exception of 1 mouse imaged on day 5), intermediate (39.5 ± 0.5 days), or late (99 ± 4 days) time points (mean ± SD). Mice were selected for imaging at random with no external markings on the cage or animal to identify whether the animal was a sham or SAH animal. Any subarachnoid blood that reached the dorsal surface of the brain during injection has cleared by post-procedure day 4 (Figure 1A). For each imaging session, mice were anesthetized with Avertin, which provides excellent RSFC signals [12]. Injectable Avertin was prepared by mixing 0.5 mL stock solution (10 g 2,2,2-tribromoethanol in 6.25 mL tert-amyl alcohol; Sigma-Aldrich, T48402-25G, and Fisher Scientific, A730-1, respectively) with 39.5 mL 0.9% normal saline, stored at 4°C for no longer than 1 month, protecting from light at all stages. To ensure a smooth induction, Avertin was injected in 2-3 divided doses intraperitoneally (0.3-0.4 mL initial dose, 0.1 mL at approximately 8 minutes, and 0.1 mL at approximately 12 minutes). More Avertin was injected if the mouse did not achieve an adequate level of anesthesia to suppress withdrawal to toe pinch. The animal was placed on a homeothermic heating pad (37.0 ± 0.1 °C) and head-fixed in a stereotaxic frame. The glass coverslip was cleaned with cotton-tipped applicators and diluted ethanol solution. Image acquisition typically began at 18-20 minutes after the initial dose of Avertin. With the exception of 4 animals, all imaging windows remained intact for the full 3-month study.

**Figure 1:**
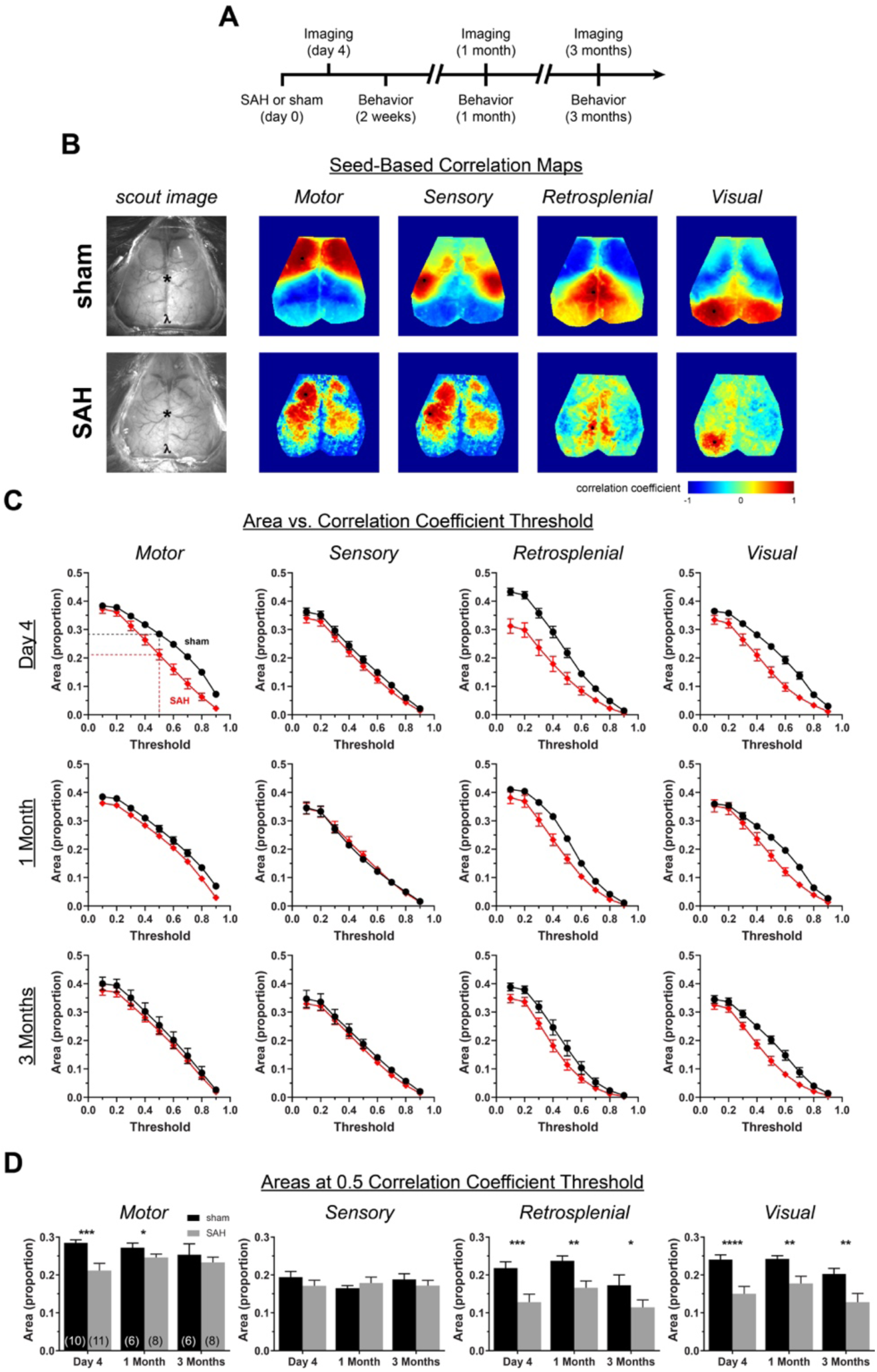
Subarachnoid hemorrhage (SAH) leads to early and late alterations in seed-based functional connectivity. (A) Experimental timeline for SAH injection, imaging, and behavioral assessments. (B) Representative seed-based connectivity maps at day 4 showing correlation coefficients from seeds placed in motor, sensory, retrosplenial, and visual cortex. Only left-sided seeds (black circles) are shown. (C) The areas above a positive seed-based correlation coefficient versus threshold are plotted at day 4, 1 month, and 3 months. Areas in proportions, normalized to total imaged brain area. (D) The proportion of total area over a correlation coefficient of 0.5. * p <0.05, ** p<0.01, *** p<0.001, **** p<0.0001 for effects between sham and SAH.

The acquisition and processing of single wavelength functional optical intrinsic signal imaging has been described previously [12, 13, 14]. The imaging surface was illuminated with a quartz tungsten halogen lamp (Techniquip R150, Capra Optical, Natick, MA) filtered at 570 ± 10 nm and directed with a fiber optic cable. μManager software was used for image acquisition [15]. Images were acquired with a Cascade 512F camera (Photometrics) at 512 x 512 pixel resolution, 11.1 FPS, and an exposure time of 50 msec for 12 minutes. A custom script was written in MATLAB (Math Works, Natick, MA, USA) for image processing. The image was down-sampled to 256 x 256 pixels. The optical density of each pixel over time was calculated and the density maps filtered between 0.008 Hz and 0.09 Hz. The signal was downsampled to 1 FPS. A brain mask was selected. The global signal was determined by taking the mean optical density for each frame. This global signal was then regressed from the optical density maps frame-by-frame over time.

Imaging data were analyzed in a blinded manner. Seed-based connectivity maps were created by mapping the correlation coefficients between seeds placed in motor, somatosensory, retrosplenial, and visual cortex and the rest of the brain. The seed locations were primarily chosen for their anatomical distribution rather than an assumed functional role. Seed coordinates were guided by the Paxinos and Franklin mouse atlas [16] and an atlas overlay adapted by White *et al.* [13] (Figure 1C). Seed-to-seed connection matrices, global connectivity maps, and interhemispheric homotopic connectivity maps and indices were calculated as described previously [12]. The connection matrix data were also organized as topological circle plots.

### Behavior

Behavioral assessments were performed at early (1-2 weeks), intermediate (approximately 1 month), and late (approximately 3 months) time points (Figure 1A). The 1 month time point was added after the study was started and, therefore, was not assessed in 6 mice. Early Y-maze and rotarod was performed at 10 ± 1 days, intermediate at 32 ± 4 days, and late at 92 ± 6 days. The first (early) MWM assessment was at 12 ± 3 days, the first reversal assessment (intermediate) was at 32 ± 4 days, and the final (late) assessment was at 93 ± 6 days. The early open field test (OFT) was performed at 13 ± 3 days, intermediate at 35 ± 3 days, and late at 95 ± 5 days (mean ± SD).

AnyMaze software (ver. 8.42, Stoelting, Wood Dale, IL, USA) was used for tracking and analysis. For the OFT, mice were placed in a 28 x 18 cm open field. Distance traveled and speed were recorded for 30 minutes. For the Y-maze, we used a Y-shaped apparatus consisting of three 33-cm arms with 15-cm-high walls, and a 7.6 x 7.6 x 7.6-cm triangular intersection. Each arm was identified with a symbol (square, circle, star). Mice were allowed access to the three arms for a total of five minutes, and their movements were recorded by AnyMaze. The number of times the mouse entered all three arms without re-entering the previous arm (i.e. triplets of ABC, ACB, BCA, etc. vs. ABA, CBC, etc.) and the total number of arm entries were recorded. From this, a percent alternation was calculated. The apparatus was cleaned with 70% ethanol between trials. For the MWM, hidden platform testing was performed in 7 trials over 3 days (trials 1-2 on day 1, trials 3-5 on day 2, and trials 6-7 on day 3). A 30-second probe test was performed 24 hours after the final hidden platform trial. Finally, a visible platform test was performed in 2 trials on the final day of testing. For the rotarod, 5 trials were performed each day for 3 days. Mice were placed on a rod at a starting rotation of 4 RPM which was constantly accelerated to 40 RPM over the course of 120 seconds. The latency of falling off of the rod and the rod RPMs were recorded.

### Statistics

Two-way repeated measures ANOVA and paired t-tests were performed with Prism 8 (GraphPad Software, Inc., CA, USA). SAS (Version 9.4, SAS Institute, Cary, NC) was used to create multivariable general linear mixed effects models. One set of models were used to determine the area (as a proportion of the imaged brain) above certain correlation coefficient thresholds (from 0.1 – 0.9) as a dependent variable of (a) condition (sham vs. SAH), (b) correlation coefficient threshold, (c) time point, and (d) region (motor, sensory, retrosplenial, or visual cortex). Hemisphere (right versus left) was assessed but did not have an effect so was removed from the final model. We also built a model to assess predictors for Y-maze percent alternation as a dependent variable of (a) condition and (b) time point. We used a p-value of 0.05 as the threshold for statistical significance. Error bars for figures are in SEM unless otherwise specified.

## RESULTS

In sham-operated mice, seed-based connectivity (i.e. correlation coefficient) maps were consistently normal when imaged longitudinally in the same animal over 3 months (Figure 1). In contrast, the SAH group showed early and persistent alterations in seed-based functional connectivity, both qualitatively and quantitatively. We found an early decrease (day 4) in motor, retrosplenial and visual cortex seed-based connectivity after SAH compared with sham, whereas sensory cortex was largely unaffected (Figure 1C, D). This decrease persisted for at least 3 months in the retrosplenial and visual seed-based maps but resolved in the motor map between 1 and 3 months. Of note, 7 of 11 mice in the SAH group showed an altered spatial pattern of connectivity and diminished correlation coefficients in the acute phase. The remaining 4 mice showed seed-based maps that were indistinguishable from sham controls (Supplemental Figure 1), suggesting a dichotomy possibly reflecting SAH severity.

As a seed-independent approach to measure functional connectivity, we determined global and interhemispheric measures of RSFC (Figure 2). There was an overall decrease in the connectivity of each pixel to every other pixel in the global connectivity map following SAH. There was also a decrease in the correlation coefficient between each pixel and its mirror pixel in contralateral hemisphere, apparent on the interhemispheric homotopic connectivity map (Figure 2A). We quantified differences between sham and SAH at different timepoints by calculating the proportional area of the imaged brain surface above correlation coefficient thresholds from 0.1 – 0.9 (Figure 2B). A decrement in global connectivity and interhemispheric connectivity following SAH is apparent at day 4 and 1 month but has largely resolved by 3 months. We also quantified global and interhemispheric connectivity observations with a Global Connectivity Index (GCI) and an Interhemispheric Connectivity Index (ICI) and found decreased GCI and ICI after SAH at early (day 4) and intermediate (1 month) time points (Figure 2C). We also noticed an interesting feature within the sham groups for GCI and IHI whereby there were differences between early and late time points (p = 0.014 and p = 0.015 for GCI and IHI with Tukey’s correction for multiple comparisons). This time course relationship was not present within the SAH group. The findings are consistent with baseline decrement of GCI and IHI under control conditions. Therefore, steady GCI and IHI over time in the SAH group might be interpreted as a relative, gradual recovery to higher connectivity values.

**Figure 2:**
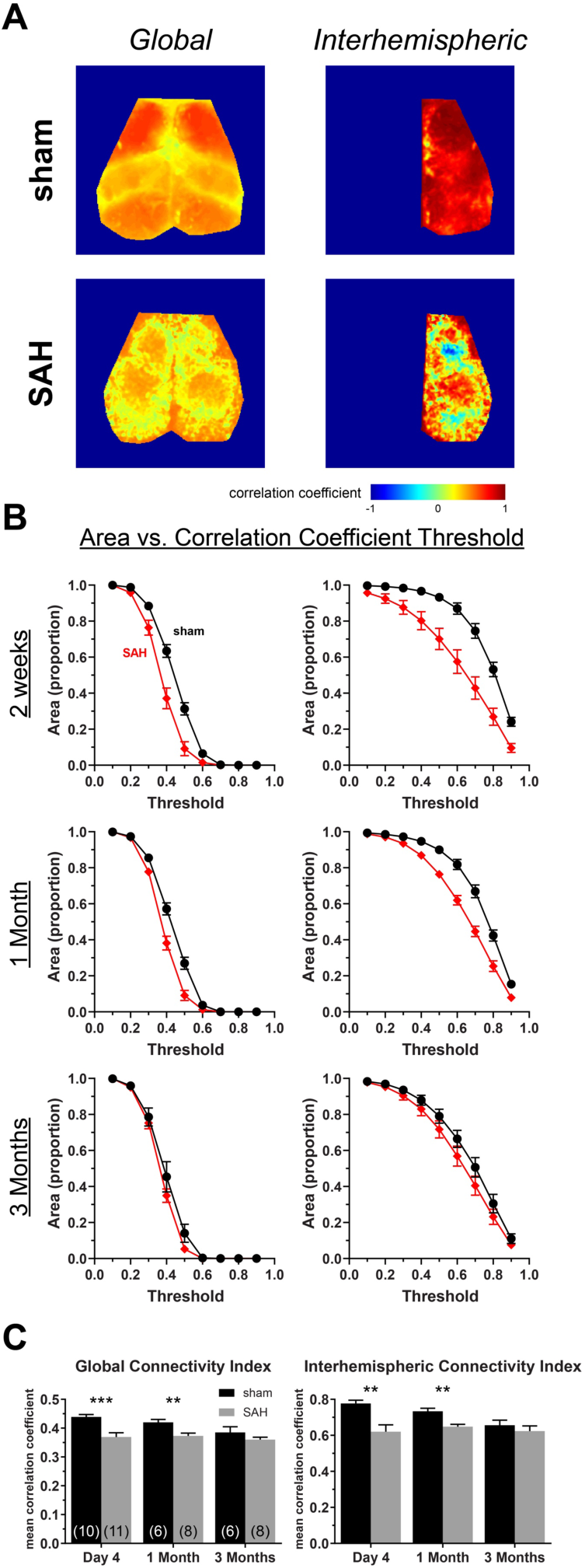
Subarachnoid hemorrhage (SAH) leads to early and intermediate changes in global and interhemispheric measures of functional connectivity. (A) Representative maps for global and interhemispheric functional connectivity in sham and SAH mice at day 4. (B) Areas are plotted as the proportion of pixels on the dorsal brain surface above a particular threshold value for global and interhemispheric maps. (C) The Global Connectivity Index and the Interhemispheric Connectivity Index at multiple time points. Number of animals for each group shown in parentheses. ** p<0.01 and *** p<0.001 for effects between sham and SAH.

We next examined the network relationship among all the seeds by generating seed-to-seed connection matrices and topological circle plots (Figure 3). As expected, we found strong interhemispheric (i.e. right vs. left) positive correlations between homotopic seeds in all four regions in sham-operated mice (Figure 3A). Weaker interhemispheric and intrahemispheric positive correlations were present between the anterior seed pairs (i.e. motor and sensory) and between the posterior seed pairs (i.e. retrosplenial and visual). Anterior seeds always appeared negatively correlated with the posterior seeds. SAH led to an early reduction in the strength of correlations, most conspicuously between the posterior seed pairs, which persisted for at least 3 months (Figure 3B, C).

**Figure 3:**
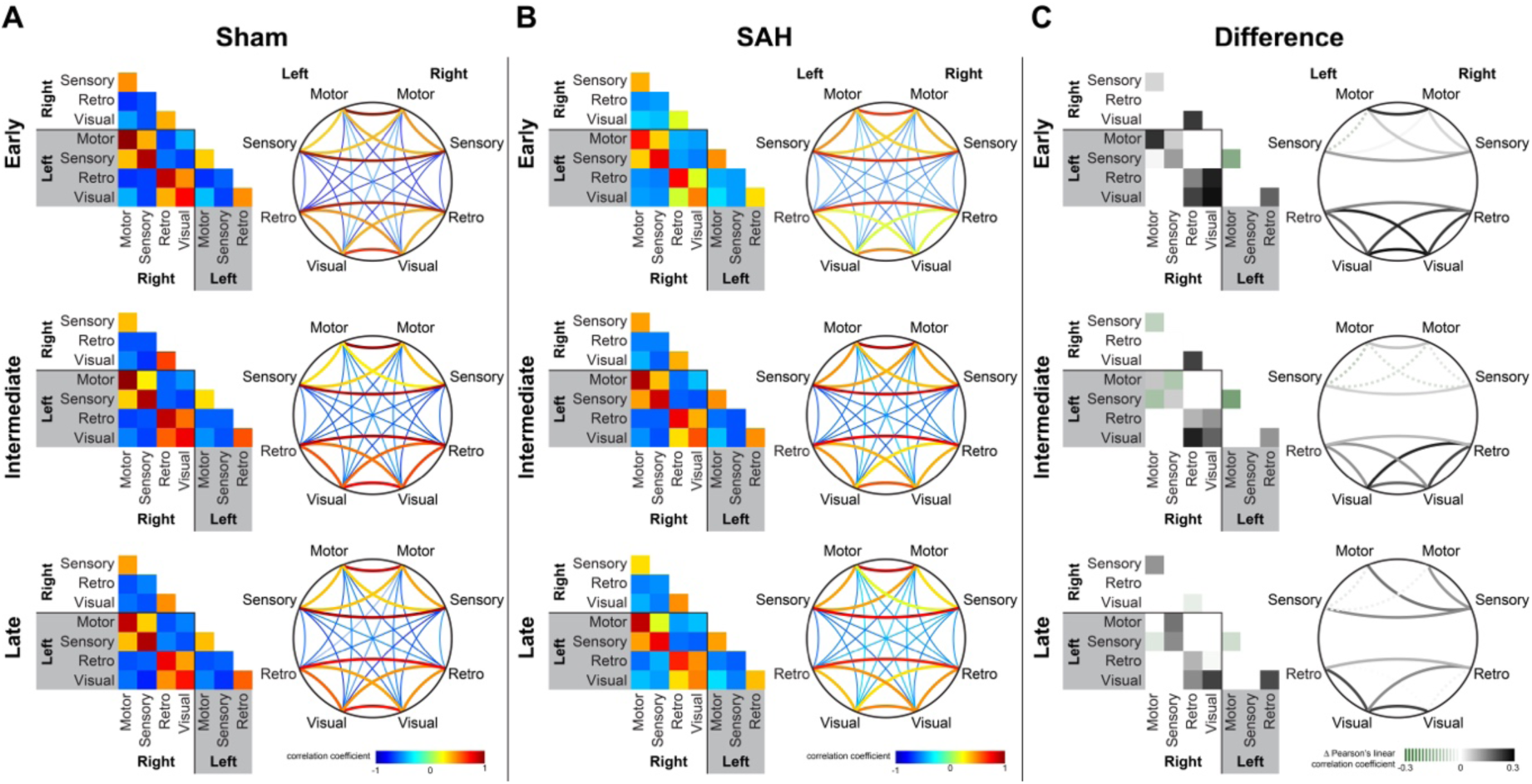
Subarachnoid hemorrhage (SAH) leads to persistent reduction in predominantly posterior functional connections. Averaged connection matrices demonstrate individual seed-to-seed correlation coefficients between motor, sensory, retrosplenial, and visual cortex in sham and SAH mice. The same data are organized as a square matrix and as a topological circle diagram for (A) sham, (B) SAH, and (C) the difference between the two at early, intermediate, and late timepoints. The difference in the connection matrix is calculated by subtracting the averaged correlation coefficient of SAH for a particular positively-correlated seed pair from that of sham. Dotted lines in (C) represent cases where SAH has a greater correlation coefficient than sham whereas solid lines in (C) represent differences where SAH has a smaller correlation coefficient than sham. Number of animals for each condition is the same as for Figure 1.

Finally, we examined the effect of SAH on the global regressed components of the optical signal fluctuations from which correlation coefficients are derived, akin to the fMRI BOLD signal (Figure 4). SAH attenuated the power of the globally regressed (i.e. hemodynamic) signal fluctuations at day 4 and to a lesser extent 1 month. In contrast, the global signal was unchanged at all time points.

**Figure 4:**
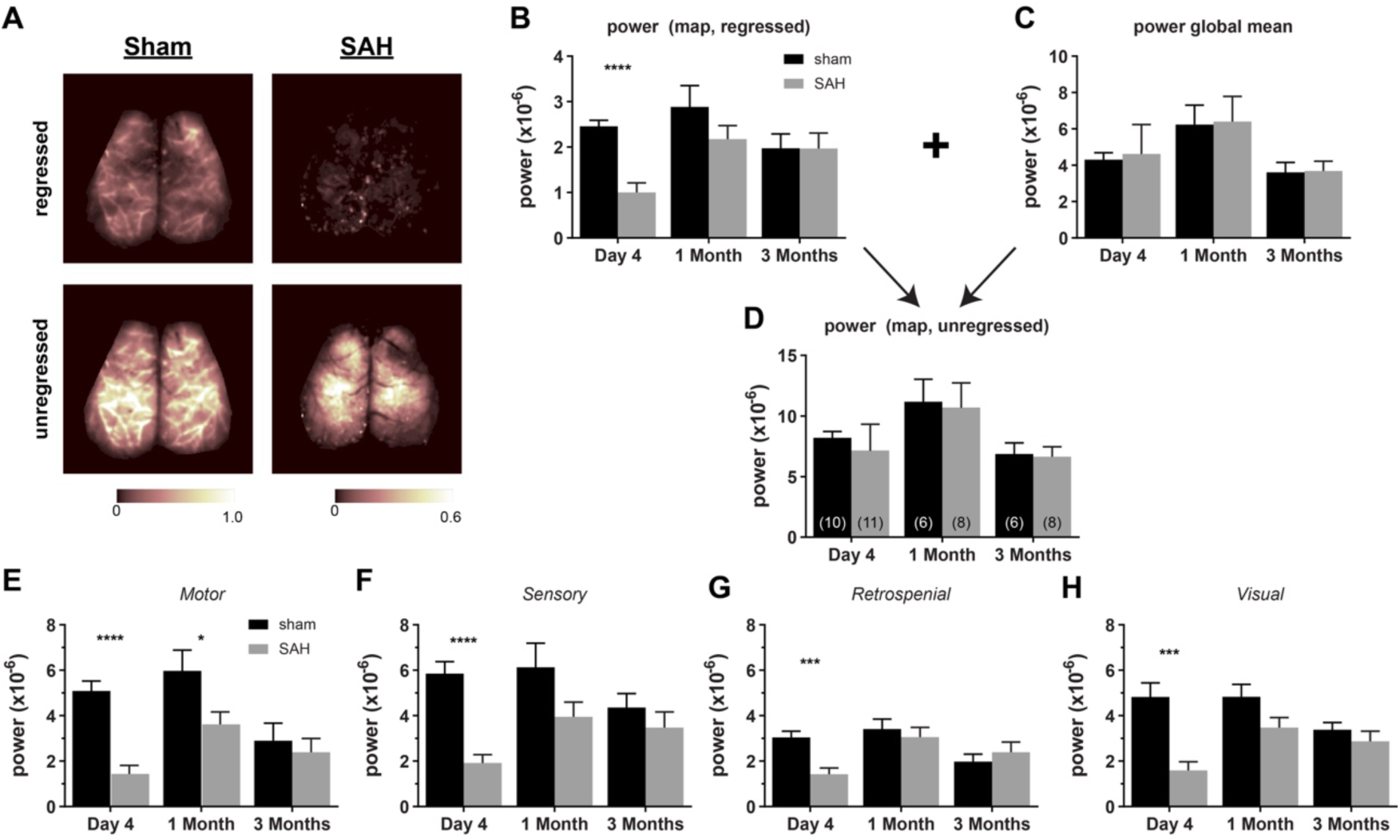
Subarachnoid hemorrhage (SAH) attenuates the amplitude of global signal regressed hemodynamic fluctuations at predominantly early time points. (A) Representative images from global signal regressed and unregressed power maps in sham and SAH mice at day 4. The power within 0.008-0.09 Hz of each pixel in the map was calculated from optical density fluctuations over the length of an imaging session. (B) Unregressed mean map powers for all mice assessed at day 4, 1 month, and 3 month time points. (C) The power of the global signal from each frame averaged over all frames with each imaging time point. (D) Global signal regressed mean map powers for all mice assessed at day 4, 1 month, and 3 month time points. (E-H) Regressed mean powers from seeds taken in (E) motor, (F) sensory, (G) retrosplenial, and (H) visual cortex. Number of animals for each group shown in parentheses. * p <0.05, *** p<0.001, **** p<0.0001 for effects between sham and SAH.

To facilitate translation of RSFC measures to potentially clinically-relevant outcomes, we performed neurocognitive assessments at early (approximately 2 weeks), intermediate (1 month), and late (3 months) time points in the imaged cohorts. SAH led to deficits on MWM hidden platform test at the early time point (Figure 5A). We could not fully assess MWM reversal at 1 month due to a disproportionate number of mice with SAH that could not swim at that time point (Supplemental Figure 2). There was 1 mouse that could swim at the early time point that could no longer swim by 1 month. We also found that at 3 months, SAH mice appeared to fatigue on the hidden platform test as trials progressed (Supplemental Figure 2). The open field test (OFT) revealed decreased distance traveled and decreased speed for SAH animals at early but not later time points (Figure 5C). However, this was not due to reduced general motor function since the Rotarod test did not reveal differences between sham and SAH mice at early (Figure 5B) and late time points (Supplemental Figure 2). The Y-maze demonstrated decreased percent alternation for SAH mice at later time points. At the early time point many of the tested mice did not meet the minimum number of entries needed (greater than 9) to adequately calculate percent alternation, rendering it uninterpretable (Figure 5D); the latter was consistent with reduced distance travelled in the open field test. We, therefore, used a mixed effects model for repeated measures at 1 and 3 months when the majority of animals could complete the test and found worse outcomes on the Y-maze after SAH compared with sham (p = 0.028, Figure 5D).

**Figure 5:**
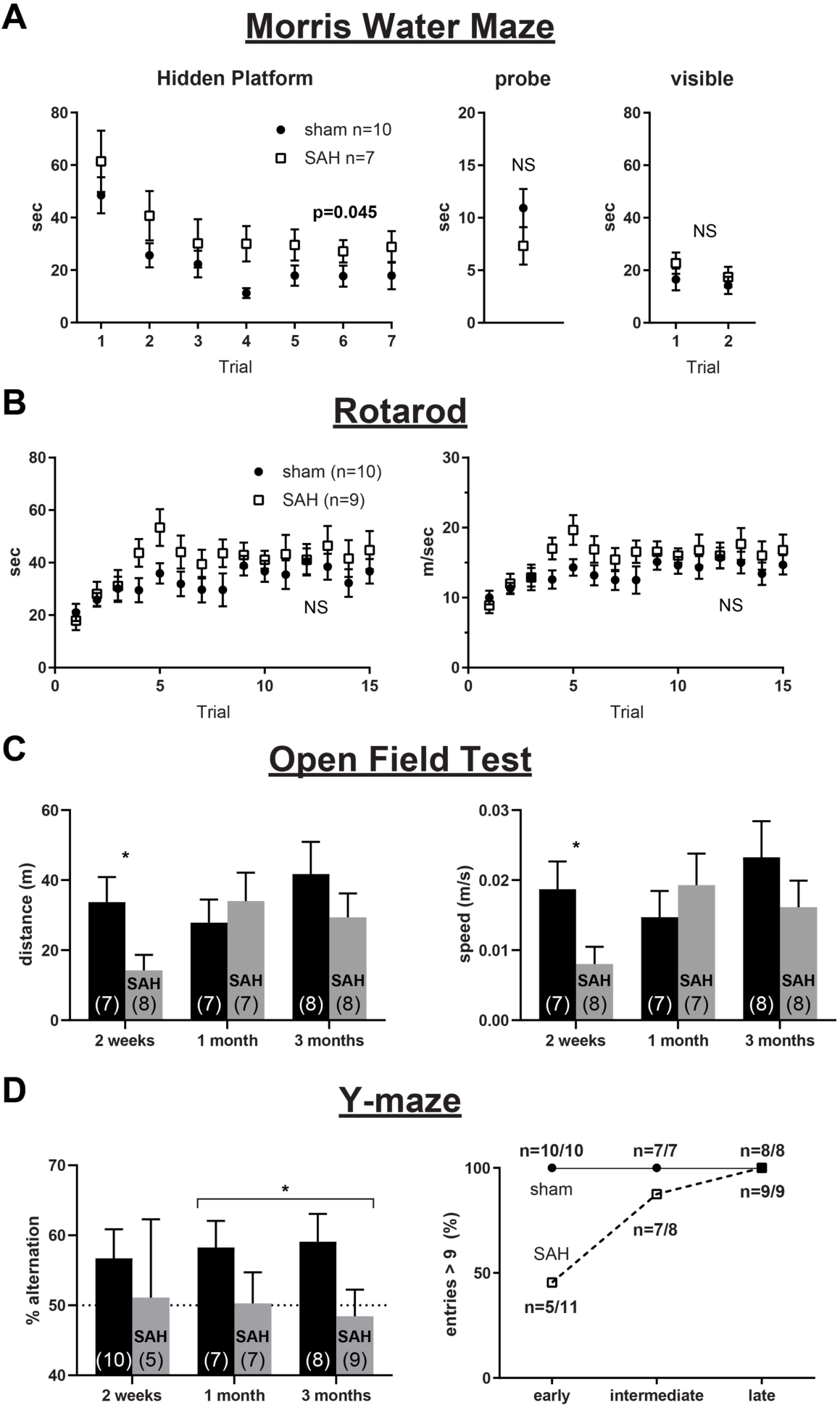
Subarachnoid hemorrhage (SAH) leads to deficits in the Morris water maze (MWM), open field test (OFT), and Y-maze. (A) Morris water maze hidden platform, probe, and visible platform performance in sham and SAH mice at approximately 2 weeks. See Supplemental Figure 2 for reversal data at 1 and 3 months. (B) Rotarod testing at 2 weeks. See Supplemental Figure 2 for similar results at 1 and 3 months. (C) OFT distance traveled and speed at 2 weeks, 1 month, and 3 month time points in sham and SAH mice. (D) Y-maze percent alternation and percent of entries greater than 9 at 2 weeks, 1 month, and 3 months in sham and SAH mice. Number of animals for each group shown in parentheses. * p <0.05

## DISCUSSION

Our data show that SAH leads to early and persistent changes in optical hemodynamic measures of resting state functional connectivity (RSFC). These changes correspond to deficits on the MWM and Y-maze. At earlier time points, there is a more general effect on imaging and behavioral findings. There also appears to be a stratification in SAH injury severity that is apparent on qualitative assessment of seed-based correlation coefficient maps. At later time points, there is persistent attenuation of RSFC measures which predominantly involves posterior network nodes. Taken together, we demonstrate that SAH causes detectable changes in blood volume-derived measures of RSFC in a well-defined mouse model of the disease.

This study employed recent advances in the optical assessment of RSFC. While fMRI is commonly used in human studies, it remains quite challenging to implement in mice [17, 18]. Therefore, White *et al.* developed a practical, minimally-invasive, optical measure of RSFC using visible light in mouse [13] which has allowed for the resolution, consistency, and throughput necessary to perform studies that assess the impact of conditions such as ischemic stroke and Alzheimer’s disease on measures of functional connectivity [19, 20]. When coupled with a chronic glass coverslip implanted over intact and unaltered skull, the technique can be a powerful way to assess RSFC longitudinally in the same animal over weeks to months [11]. Another feature of the method is that it assesses blood volume fluctuations that are akin to the BOLD signal utilized in fMRI studies [18, 21]. Therefore, observations using the optical approach may be comparable to fMRI studies.

The specific behavioral findings that we report are novel, but other early and late behavioral deficits have been described previously by others. We find deficits in the MWM at 2 weeks, and it is worth noting that our MWM assessments at 1 month and 3 months are reversal tests (i.e. all mice are influenced by the earlier 2 week MWM testing). Others have described deficits in the MWM following SAH in mice using different experimental timelines. Using the same injection approach that we used, one group performed the MWM prior to SAH induction and found reversal of performance in SAH mice assessed for up to 7 days [22, 23]. Interestingly, Provencio *et al.*, used a unique model of venous SAH which does not reproduce acute elevations in ICP and found that SAH leads to deficits on the first assessment of the MWM at 1 month [24]. They also used the modified Barnes maze with results that might suggest early and delayed cognitive deficits [25, 24]. Therefore, although our RSFC imaging findings could be considered an early brain injury biomarker, our behavioral tests at approximately 2 weeks and beyond do not distinguish between early and delayed injury. We are not aware of other reports of Y-maze testing in the anterior prechiasmatic injection model of SAH. We found that it was useful to have the open field test (OFT) as an independent assessment of ambulation which strengthened our impression that early assessment of the Y-maze was confounded by decreased ambulation. As the ambulation and activity measures resolved, we were able to detect a signal for persistent SAH-associated deficits of percent alternation on the Y-maze. Interestingly, we found no deficits on the rotarod whereas prior studies using an endovascular perforation model of SAH found early deficits [26–28]. The difference from our findings could be due to earlier outcome assessment in the other studies (which were at day 3) or due to effects of early focal ischemic stroke that is associated with the endovascular perforation model. For our purposes, we view negative rotarod testing as a helpful control that strengthens the specificity of our findings for predominant deficits in spatial learning and working memory.

We did not assess for structural or anatomical changes to explain persistent long-term deficits in RSFC or behavior at 3 months. Prior work observed diffuse neuronal death up to 7 days after SAH induction using the anterior prechiasmatic injection approach used for the current study [9, 29, 23, 22]. Additional studies using the endovascular perforation model have reported acute white matter injury—independent of ischemic stroke—using MRI [30–32], ultrastructural assessment with electron microscopy [32], and assessment of molecular markers for white matter injury, including beta-APP [30]. Notably, investigators have described posterior dominant involvement of white matter tracts following an endovascular puncture model of SAH [33] which is consistent with our findings of persistent posterior-dominant deficits in RSFC (Figures 1 and 3). Future RSFC studies designed to look specifically at markers of neuronal damage and location of white matter tracts at both early and late time points in additional models of SAH would be informative.

Another question raised by our findings is if persistent changes in RSFC or behavior are due to changes in neuronal activity, problems with neurovascular coupling, or both. We assessed a hemodynamic measure of RSFC. We took this approach because it provides the most accessible means to study RSFC in mouse and is clinically analogous to the fMRI BOLD signal used in humans. However, prior animal studies suggest that there can be neuronal hyperexcitability [34] rather than a depression of activity. There is also a body of evidence suggesting that neurovascular coupling is diminished following acute brain injury, including SAH [35]; however, the exact time course of neurovascular changes has not been fully defined. Regardless of underlying mechanisms, we have described what we hope to be, at a minimum, a useful biomarker for early brain injury after SAH. Future studies using genetically-encoded neuronal calcium fluorescence reporters (i.e. GCaMP mice) or invasive electrophysiology [36–39] might provide additional insight into what underlies changes in our observed hemodynamically-derived measures of RSFC.

It is worth commenting on our choice of a mouse model. There are several mouse models of subarachnoid hemorrhage [40, 41]; however, a perfect SAH model does not exist and there is not yet a consensus about which model to use for each experimental condition [42]. We chose to base our study on the anterior prechiasmatic injection model which recapitulates early brain injury, leads to early elevations in ICP, and results in a significant clot burden at the base of the skull [9]. Another approach we considered was the endovascular perforation model wherein a filament is advanced into the middle cerebral artery until the vessel is perforated [43]. While the endovascular puncture model remains an essential approach in the field, we aimed to avoid the acute focal cerebral infarcts that can occur with this method. Injection of arterial blood through the cisternal magna is another accepted model; however, it does not fully reproduce the persistent elevated ICP that is seen with aneurysm rupture. Similarly, transection of a subarachnoid vein overlying the cisterna magna is another approach which further isolates effects of blood itself in the pathogenesis of SAH, but again does not lead to the acute ICP elevations desirable for an early brain injury model [24]. Finally, the spontaneous aneurysm rupture approach could be seen as the most clinically analogous mouse model, but variability and the unpredictability of when a rupture occurs precluded its use in the current study. It is worth emphasizing that now that we have provided a description of our findings in one model of SAH, we hope that measures of functional connectivity will be tested in additional mouse models and strains. Indeed, since there is no one perfect model of SAH, we believe that the only way to approach a useful understanding of the mechanisms and clinically relevant outcomes involved is to attempt similar investigations using multiple different models in different laboratories.

Our study has additional limitations. We measured RSFC under anesthesia. We did this because the meaning and feasibility of optical measures of RSFC in awake animals has been thrown into question [44]. Therefore, we recently performed a systematic study to determine the optimal anesthetic for our purposes and found that Avertin (tribromoethanol) anesthesia provided the most robust results [12]. Regardless, our findings would be strengthened if future studies attempted awake assessments or assessments under other types of anesthesia. Also, we do not draw conclusions about why seeds with altered connectivity (in retrosplenial cortex, for example) are associated with the observed behavioral findings. Finally, we only used male mice in our study. Future studies should use both male and female mice.

In conclusion, we find that SAH leads to early and persistent changes in resting state functional connectivity and deficits in spatial learning and working memory tasks. At a minimum, we have demonstrated how optical measures of hemodynamically-derived functional connectivity could serve as a non-invasive and longitudinal biomarker for severity in experimental SAH. We hope that our observations will serve to guide future work to determine mechanisms potentially linking SAH to altered functional connectivity and, ultimately, to better understand how functional connectivity might affect cognitive outcomes following SAH.

## ACKNOWLEDGEMENTS

We thank Jinglu Ai, Hoyee Wan, and the Macdonald Lab for helpful advice regarding the anterior prechiasmatic injection mouse model. Additional thanks to Brian Edlow, Leigh Hochberg, and Aman Patel for useful discussions about the connectivity data and to James Chung for comments on the manuscript.

## REFERENCES

1. Bederson JB, Connolly ES, Jr., Batjer HH, Dacey RG, Dion JE, Diringer MN et al. Guidelines for the management of aneurysmal subarachnoid hemorrhage: a statement for healthcare professionals from a special writing group of the Stroke Council, American Heart Association. Stroke; a journal of cerebral circulation. 2009;40(3):994–1025. doi:10.1161/STROKEAHA.108.191395.

2. Taylor TN, Davis PH, Torner JC, Holmes J, Meyer JW, Jacobson MF. Lifetime cost of stroke in the United States. Stroke; a journal of cerebral circulation. 1996;27(9):1459–66.

3. Ogden JA, Mee EW, Henning M. A prospective study of impairment of cognition and memory and recovery after subarachnoid hemorrhage. Neurosurgery. 1993;33(4):572–86; discussion 86-7.

4. Passier PE, Visser-Meily JM, Rinkel GJ, Lindeman E, Post MW. Life satisfaction and return to work after aneurysmal subarachnoid hemorrhage. Journal of stroke and cerebrovascular diseases: the official journal of National Stroke Association. 2011;20(4):324–9. doi:10.1016/j.jstrokecerebrovasdis.2010.02.001.

5. Ellmore TM, Rohlffs F, Khursheed F. FMRI of working memory impairment after recovery from subarachnoid hemorrhage. Front Neurol. 2013;4:179. doi:10.3389/fneur.2013.00179.

6. Maher M, Churchill NW, de Oliveira Manoel AL, Graham SJ, Macdonald RL, Schweizer TA. Altered Resting-State Connectivity within Executive Networks after Aneurysmal Subarachnoid Hemorrhage. PloS one. 2015;10(7):e0130483. doi:10.1371/journal.pone.0130483.

7. da Costa L, Dunkley BT, Bethune A, Robertson A, Keller A, Pang EW. Increased Frontal Lobe Activation After Aneurysmal Subarachnoid Hemorrhage. Stroke; a journal of cerebral circulation. 2016. doi:10.1161/STROKEAHA.116.013786.

8. Kilkenny C, Browne WJ, Cuthill IC, Emerson M, Altman DG. Improving bioscience research reporting: the ARRIVE guidelines for reporting animal research. PLoS Biol. 2010;8(6):e1000412. doi:10.1371/journal.pbio.1000412.

9. Sabri M, Jeon H, Ai J, Tariq A, Shang X, Chen G et al. Anterior circulation mouse model of subarachnoid hemorrhage. Brain research. 2009;1295:179–85. doi:10.1016/j.brainres.2009.08.021.

10. Oka F, Hoffmann U, Lee JH, Shin HK, Chung DY, Yuzawa I et al. Requisite ischemia for spreading depolarization occurrence after subarachnoid hemorrhage in rodents. Journal of cerebral blood flow and metabolism: official journal of the International Society of Cerebral Blood Flow and Metabolism. 2016. doi:10.1177/0271678X16659303.

11. Silasi G, Xiao D, Vanni MP, Chen AC, Murphy TH. Intact skull chronic windows for mesoscopic wide-field imaging in awake mice. Journal of neuroscience methods. 2016;267:141–9. doi:10.1016/j.jneumeth.2016.04.012.

12. Xie H, Chung DY, Kura S, Sugimoto K, Aykan SA, Wu Y et al. Differential effects of anesthetics on resting state functional connectivity in the mouse. Journal of cerebral blood flow and metabolism: official journal of the International Society of Cerebral Blood Flow and Metabolism. 2019:271678X19847123. doi:10.1177/0271678X19847123.

13. White BR, Bauer AQ, Snyder AZ, Schlaggar BL, Lee JM, Culver JP. Imaging of functional connectivity in the mouse brain. PloS one. 2011;6(1):e16322. doi:10.1371/journal.pone.0016322.

14. Kura S, Xie H, Fu B, Ayata C, Boas DA, Sakadzic S. Intrinsic optical signal imaging of the blood volume changes is sufficient for mapping the resting state functional connectivity in the rodent cortex. J Neural Eng. 2018. doi:10.1088/1741-2552/aaafe4.

15. Edelstein AD, Tsuchida MA, Amodaj N, Pinkard H, Vale RD, Stuurman N. Advanced methods of microscope control using muManager software. J Biol Methods. 2014;1(2). doi:10.14440/jbm.2014.36.

16. Paxinos G, Franklin KBJ. The mouse brain in stereotaxic coordinates. 2. ed. San Diego: Academic Press; 2001.

17. Pan WJ, Billings JC, Grooms JK, Shakil S, Keilholz SD. Considerations for resting state functional MRI and functional connectivity studies in rodents. Front Neurosci. 2015;9:269. doi:10.3389/fnins.2015.00269.

18. Desjardins M, Kılıç K, Thunemann M, Mateo C, Holland D, Ferri CGL et al. Awake mouse imaging: from 2-photon microscopy to BOLD fMRI. Biol Psychiatry Cogn Neurosci Neuroimaging. 2019.

19. Bauer AQ, Kraft AW, Wright PW, Snyder AZ, Lee JM, Culver JP. Optical imaging of disrupted functional connectivity following ischemic stroke in mice. NeuroImage. 2014;99:388–401. doi:10.1016/j.neuroimage.2014.05.051.

20. Bero AW, Bauer AQ, Stewart FR, White BR, Cirrito JR, Raichle ME et al. Bidirectional relationship between functional connectivity and amyloid-beta deposition in mouse brain. The Journal of neuroscience: the official journal of the Society for Neuroscience. 2012;32(13):4334–40. doi:10.1523/JNEUROSCI.5845-11.2012.

21. Hillman EM. Coupling mechanism and significance of the BOLD signal: a status report. Annual review of neuroscience. 2014;37:161–81. doi:10.1146/annurevneuro-071013-014111.

22. Schallner N, Pandit R, LeBlanc R, 3rd, Thomas AJ, Ogilvy CS, Zuckerbraun BS et al. Microglia regulate blood clearance in subarachnoid hemorrhage by heme oxygenase-1. The Journal of clinical investigation. 2015;125(7):2609–25. doi:10.1172/JCI78443.

23. LeBlanc RH, 3rd, Chen R, Selim MH, Hanafy KA. Heme oxygenase-1-mediated neuroprotection in subarachnoid hemorrhage via intracerebroventricular deferoxamine. J Neuroinflammation. 2016;13(1):244. doi:10.1186/s12974-016-0709-1.

24. Provencio JJ, Swank V, Lu H, Brunet S, Baltan S, Khapre RV et al. Neutrophil depletion after subarachnoid hemorrhage improves memory via NMDA receptors. Brain Behav Immun. 2016;54:233–42. doi:10.1016/j.bbi.2016.02.007.

25. Provencio JJ, Altay T, Smithason S, Moore SK, Ransohoff RM. Depletion of Ly6G/C(+) cells ameliorates delayed cerebral vasospasm in subarachnoid hemorrhage. J Neuroimmunol. 2011;232(1-2):94–100. doi:10.1016/j.jneuroim.2010.10.016.

26. Gao J, Wang H, Sheng H, Lynch JR, Warner DS, Durham L et al. A novel apoE-derived therapeutic reduces vasospasm and improves outcome in a murine model of subarachnoid hemorrhage. Neurocrit Care. 2006;4(1):25–31. doi:10.1385/NCC:4:1:025.

27. Mesis RG, Wang H, Lombard FW, Yates R, Vitek MP, Borel CO et al. Dissociation between vasospasm and functional improvement in a murine model of subarachnoid hemorrhage. Neurosurg Focus. 2006;21(3):E4.

28. Wang H, Gao J, Lassiter TF, McDonagh DL, Sheng H, Warner DS et al. Levetiracetam is neuroprotective in murine models of closed head injury and subarachnoid hemorrhage. Neurocrit Care. 2006;5(1):71–8. doi:10.1385/NCC:5:1:71.

29. Boettinger S, Kolk F, Broessner G, Helbok R, Pfausler B, Schmutzhard E et al. Behavioral characterization of the anterior injection model of subarachnoid hemorrhage. Behav Brain Res. 2017;323:154–61. doi:10.1016/j.bbr.2017.02.004.

30. Egashira Y, Hua Y, Keep RF, Xi G. Acute white matter injury after experimental subarachnoid hemorrhage: potential role of lipocalin 2. Stroke; a journal of cerebral circulation. 2014;45(7):2141–3. doi:10.1161/STROKEAHA.114.005307.

31. Egashira Y, Zhao H, Hua Y, Keep RF, Xi G. White Matter Injury After Subarachnoid Hemorrhage: Role of Blood-Brain Barrier Disruption and Matrix Metalloproteinase-9. Stroke; a journal of cerebral circulation. 2015;46(10):2909–15. doi:10.1161/STROKEAHA.115.010351.

32. Kummer TT, Magnoni S, MacDonald CL, Dikranian K, Milner E, Sorrell J et al. Experimental subarachnoid haemorrhage results in multifocal axonal injury. Brain: a journal of neurology. 2015;138(Pt 9):2608–18. doi:10.1093/brain/awv180.

33. Wu Y, Peng J, Pang J, Sun X, Jiang Y. Potential mechanisms of white matter injury in the acute phase of experimental subarachnoid haemorrhage. Brain: a journal of neurology. 2017;140(6):e36. doi:10.1093/brain/awx084.

34. Li B, Luo C, Tang W, Chen Z, Li Q, Hu B et al. Role of HCN channels in neuronal hyperexcitability after subarachnoid hemorrhage in rats. The Journal of neuroscience: the official journal of the Society for Neuroscience. 2012;32(9):3164–75. doi:10.1523/JNEUROSCI.5143-11.2012.

35. Balbi M, Koide M, Wellman GC, Plesnila N. Inversion of neurovascular coupling after subarachnoid hemorrhage in vivo. Journal of cerebral blood flow and metabolism: official journal of the International Society of Cerebral Blood Flow and Metabolism. 2017:271678X16686595. doi:10.1177/0271678X16686595.

36. Ma Y, Shaik MA, Kim SH, Kozberg MG, Thibodeaux DN, Zhao HT et al. Wide-field optical mapping of neural activity and brain haemodynamics: considerations and novel approaches. Philos Trans R Soc Lond B Biol Sci. 2016;371(1705). doi:10.1098/rstb.2015.0360.

37. Ma Y, Shaik MA, Kozberg MG, Kim SH, Portes JP, Timerman D et al. Resting-state hemodynamics are spatiotemporally coupled to synchronized and symmetric neural activity in excitatory neurons. Proceedings of the National Academy of Sciences of the United States of America. 2016;113(52):E8463–E71. doi:10.1073/pnas.1525369113.

38. Xiao D, Vanni MP, Mitelut CC, Chan AW, LeDue JM, Xie Y et al. Mapping cortical mesoscopic networks of single spiking cortical or sub-cortical neurons. Elife. 2017;6. doi:10.7554/eLife.19976.

39. Wright PW, Brier LM, Bauer AQ, Baxter GA, Kraft AW, Reisman MD et al. Functional connectivity structure of cortical calcium dynamics in anesthetized and awake mice. PloS one. 2017;12(10):e0185759. doi:10.1371/journal.pone.0185759.

40. Jeon H, Ai J, Sabri M, Tariq A, Shang X, Chen G et al. Neurological and neurobehavioral assessment of experimental subarachnoid hemorrhage. BMC Neurosci. 2009;10:103. doi:10.1186/1471-2202-10-103.

41. Oka F, Chung DY, Suzuki M, Ayata C. Delayed Cerebral Ischemia After Subarachnoid Hemorrhage: Experimental-Clinical Disconnect and the Unmet Need. Neurocrit Care. 2019. doi:10.1007/s12028-018-0650-5.

42. Leclerc JL, Garcia JM, Diller MA, Carpenter AM, Kamat PK, Hoh BL et al. A Comparison of Pathophysiology in Humans and Rodent Models of Subarachnoid Hemorrhage. Front Mol Neurosci. 2018;11:71. doi:10.3389/fnmol.2018.00071.

43. Parra A, McGirt MJ, Sheng H, Laskowitz DT, Pearlstein RD, Warner DS. Mouse model of subarachnoid hemorrhage associated cerebral vasospasm: methodological analysis. Neurological research. 2002;24(5):510–6. doi:10.1179/016164102101200276.

44. Winder AT, Echagarruga C, Zhang Q, Drew PJ. Weak correlations between hemodynamic signals and ongoing neural activity during the resting state. Nature neuroscience. 2017;20(12):1761–9. doi:10.1038/s41593-017-0007-y.

