## Supplementary figures and images for "Subarachnoid hemorrhage leads to early and persistent functional connectivity and behavioral changes in mice"

### Supplemental Figure 1

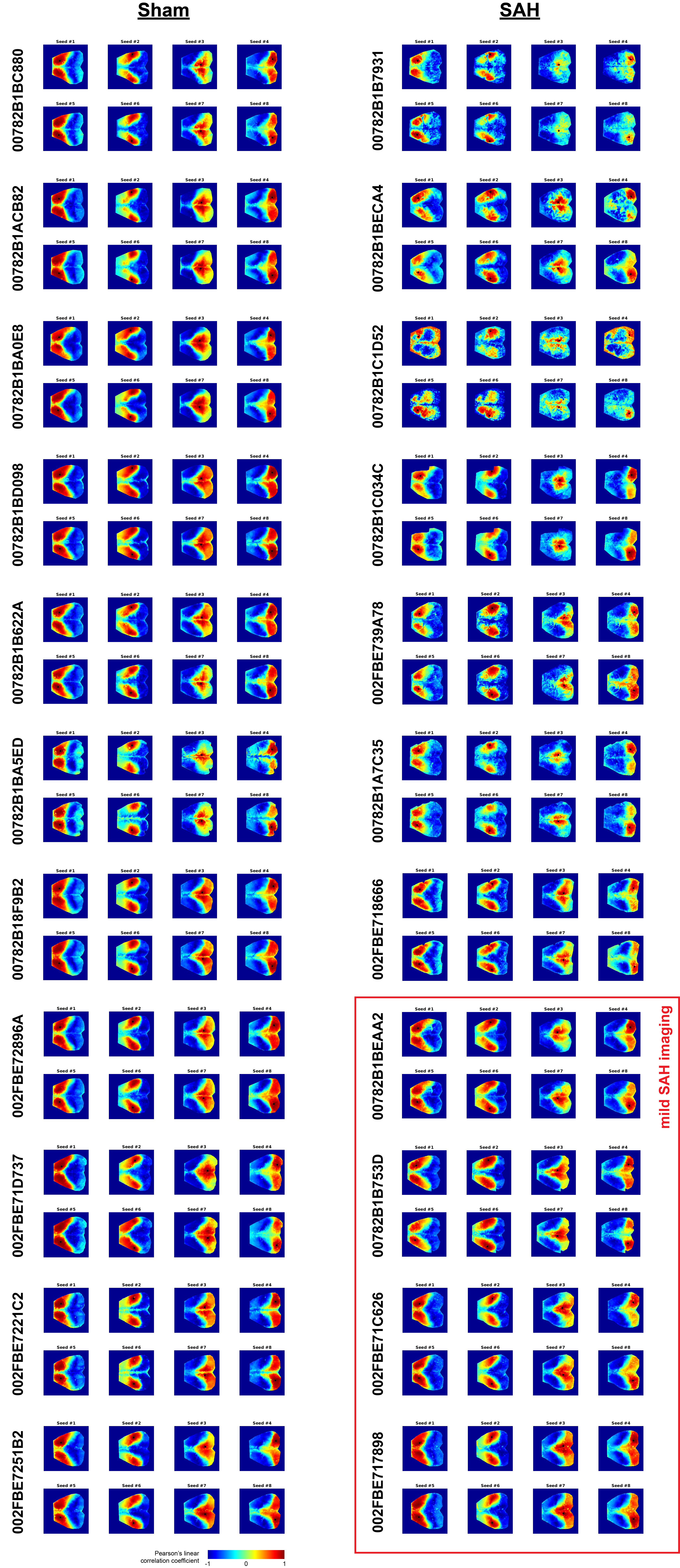

### Supplemental Figure 2

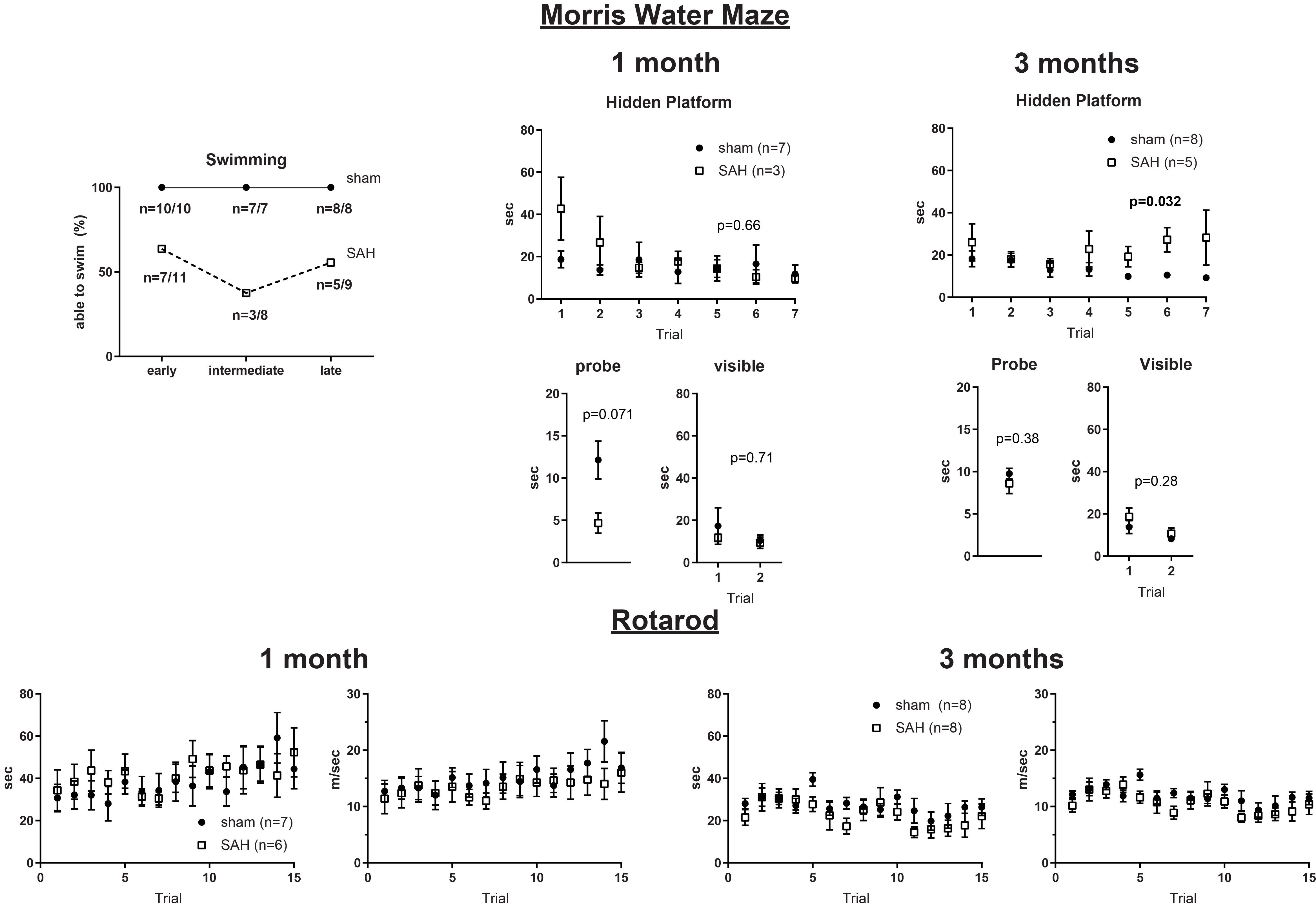
