## Supplemental Figure Captions for "Subarachnoid hemorrhage leads to early and persistent functional connectivity and behavioral changes in mice"

**Supplemental Figure 1:** Representative seed-based correlation coefficient maps for all early sham and SAH imaging. Each map is identified by its 12-digit hexadecimal radiofrequency identification tag (RFID). SAH maps in the red box were qualitatively determined to have relatively intact seed-based connectivity.

**Supplemental Figure 2:** Additional behavioral findings. The ability of the mouse in each cohort to swim is plotted for early, intermediate and late time points. MWM was only performed and analyzed for those mice which could swim. Additional hidden platform, probe, and visible platform testing at 1 and 3 months is also shown. Rotarod testing at 1 and 3 months demonstrate no differences between SAH and sham.
